# Sphingolipid changes in Parkinson L444P GBA mutation fibroblasts promote α-synuclein aggregation

**DOI:** 10.1101/2020.11.09.375048

**Authors:** Céline Galvagnion, Silvia Cerri, Anthony H.V. Schapira, Fabio Blandini, Donato A. Di Monte

**Affiliations:** German Center for Neurodegenerative Diseases (DZNE), Venusberg-Campus 1, Building 99, 53127 Bonn, Germany; Department of Drug Design and Pharmacology, Faculty of Health and Medical Sciences, University of Copenhagen, Universitetsparken 2, 2100, Copenhagen Ø, Denmark; Laboratory of Cellular and Molecular Neurobiology, IRCCS Mondino Foundation, Pavia, Italy; Department of Clinical and Movement Neurosciences, UCL Queen Square Institute of Neurology, London, UK; Department of Brain and Behavioral Sciences, University of Pavia, Pavia, Italy

**Keywords:** Fibroblasts, GBA, α-synuclein, lipidomics, Parkinson’s Disease

## Abstract

Intraneuronal accumulation of aggregated α-synuclein is a pathological hallmark of Parkinson’s disease. Therefore, mechanisms capable of promoting α-synuclein deposition bear important pathogenetic implications. Mutations of the glucocerebrosidase 1 (*GBA*) gene represent a prevalent Parkinson’s disease risk factor. They are associated with loss of activity of a key enzyme involved in lipid metabolism, glucocerebrosidase, supporting a mechanistic relationship between abnormal α-synuclein-lipid interactions and the development of Parkinson pathology. In this study, the lipid membrane composition of fibroblasts isolated from control subjects, patients with idiopathic Parkinson’s disease (iPD) and Parkinson patients carrying the L444P *GBA* mutation (PD-GBA) was assayed using shotgun lipidomics. The lipid profile of PD-GBA fibroblasts differed significantly from that of control and iPD cells. It was characterized by an overall increase in sphingolipid levels. It also featured a significant change in the proportion of ceramide, sphingomyelin and hexosylceramide molecules with shorter and longer hydrocarbon chain length; levels of shorter-chain molecules were increased while the percent of longer-chain sphingolipids was decreased in PD-GBA lipid extracts. The extent of this shift was correlated to the degree of reduction of fibroblast glucocerebrosidase activity. In a second set of experiments, lipid extracts from control and PD-GBA fibroblasts were added to incubations of recombinant α-synuclein. The kinetics of α-synuclein aggregation, as assessed by the binding of thioflavin T to amyloid structures, was significantly accelerated after addition of PD-GBA extracts as compared to control samples. Amyloid fibrils collected at the end of these incubations contained lipids, indicating α-synuclein-lipid co-assembly. Lipids extracted from α-synuclein fibrils were also analysed by shotgun lipidomics. Data revealed that the lipid content of these fibrils was significantly enriched of shorter-chain sphingolipids. Taken together, findings of this study indicate that the L444P *GBA* mutation and consequent enzymatic loss are associated with a distinctly altered membrane lipid profile that provides a biological fingerprint of this mutation in Parkinson fibroblasts. This altered lipid profile, which includes an increased content of shorter-chain sphingolipids, could also be an indicator of increased risk for α-synuclein aggregate pathology. Shorter-chain molecules may act as preferred reactants during lipid-induced α-synuclein fibrillation.

## INTRODUCTION

Mutations of the glucocerebrosidase 1 (*GBA*) gene have long been associated with Gaucher’s disease but also represent the commonest known genetic risk factor for Parkinson’s disease (Schapira, 2015; Ryan *et al*., 2019). Parkinson risk is significantly higher in mutation carriers, with variable odds ratios that depend on the specific *GBA* mutation and racial characteristics of the population tested (Zhang *et al*., 2018). Disease penetrance also varies with age and has been estimated to range from 10 to 30% in mutation carriers aged 50 to 80 years old (Anheim *et al*., 2012). Clinical features are indistinguishable between Parkinson patients carrying *GBA* mutations (PD-GBA) and patients affected by idiopathic Parkinson’s Disease (iPD), as underscored by investigations showing that a significant proportion of “iPD” patients (approximately 5-10%), once specifically tested, were in fact carriers of *GBA* mutations (Neumann *et al*., 2009; Sidransky *et al*., 2009; Alcalay *et al*., 2012; Schapira, 2015). Genotype-phenotype correlation studies have confirmed the similarities in clinical and pathological manifestations between PD-GBA and iPD. They have also revealed that PD-GBA is characterized by a slightly earlier disease onset and higher prevalence of cognitive impairment and other non-motor symptoms (Neumann *et al*., 2009; Brockmann *et al*., 2011; Winder-Rhodes *et al*., 2013). Different *GBA* mutations may be associated with different clinical phenotypes, with more severe parkinsonian features and a more aggressive disease course affecting PD-GBA patients carrying specific “severe” variants, such as the p-L444P mutation (Gan-Or *et al*., 2015; Cilia *et al*., 2016).

*GBA* encodes the lysosomal enzyme glucocerebrosidase (GCase), and *GBA* mutations cause a decrease in GCase activity and GCase-catalyzed hydrolysis of glucosylceramide (GluCer) to glucose and ceramide (Cer). Loss of GCase activity has been reported in the brain and blood of PD-GBA patients as well as in patient-derived cells, such as skin fibroblasts and dopaminergic neurons generated from induced pluripotent stem cells (iPSCs) (Gegg *et al*., 2012; Schöndorf *et al*., 2014; Alcalay *et al*., 2015; Aflaki *et al*., 2016; Sanchez-Martinez *et al*., 2016; Collins *et al*., 2018; Moors *et al*., 2019). Decreased GCase activity would be expected to result in an accumulation of GCase substrates as seen, for example, in iPSCs from PD-GBA patients containing higher levels of GluCer and glucosylsphingosine (GluSph) (Schöndorf *et al*., 2014; Aflaki *et al*., 2016). It is noteworthy, however, that due to the close interrelationship between pathways of lipid metabolism and the central role played by Cer in sphingolipid (SL) homeostasis, lower GCase activity is likely to have broader consequences on cell lipid composition and, in particular, SL synthesis, maintenance and breakdown (Fig. 1) (Futerman and Platt, 2017). To date, only limited information is available from lipidomic analyses of cell or tissue samples from PD-GBA patients (Gegg *et al*., 2012; Schöndorf *et al*., 2014; Aflaki *et al*., 2016). More detailed and comprehensive studies are, therefore, warranted for the identification of PD-GBA lipid signatures. These studies could also shed light upon the role that specific changes in lipid profile may play in increasing the risk for PD pathology.

**Figure 1.**
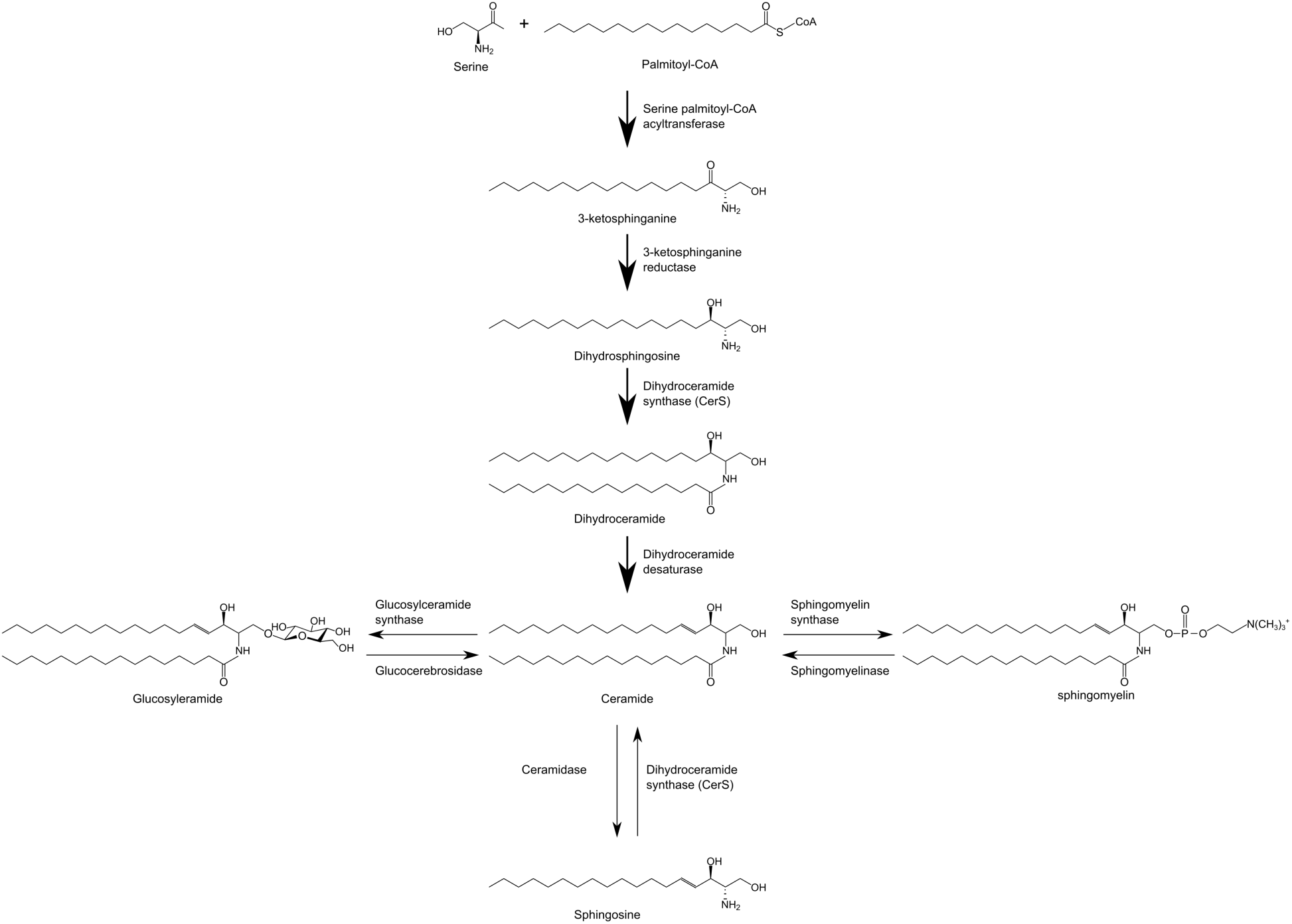
Simplified sphingolipid metabolic pathway.

Mechanisms contributing to PD pathogenesis in carriers of *GBA* mutations are not fully understood. Of likely relevance, however, is evidence indicating a reciprocal relationship between GCase activity and intracellular levels and toxic properties of α-synuclein (αS) (Mazzulli *et al*., 2011; Schapira *et al*., 2014). αS is a key player in the pathogenesis of PD. It is a major component of Lewy bodies and Lewy neurites, the intraneuronal inclusions pathognomonic of iPD (Spillantini *et al*., 1997). Furthermore, single-point and multiplication mutations of the αS gene are causally associated with familial forms of parkinsonism (Polymeropoulos *et al*., 1997; Nussbaum, 2018; Bandres-Ciga *et al*., 2020). The tendency of αS to assemble into oligomeric and fibrillar aggregates is thought to be a gain of toxic function involved in inclusion formation and other neurodegenerative effects. Interestingly, loss of GCase activity, as induced by *GBA* mutations, causes intracellular accumulation of monomeric as well as aggregated αS (Mazzulli *et al*., 2011; Schöndorf *et al*., 2014; Aflaki *et al*., 2016; Yang *et al*., 2017, 2020; Gündner *et al*., 2019). Increased αS levels are in turn capable of reducing GCase activity, giving rise to a reciprocal self-amplifying cycle of protein accumulation and enzyme inhibition (Mazzulli *et al*., 2011; Gegg *et al*., 2012; Yap *et al*., 2013; Schapira *et al*., 2014). Changes in lipid composition caused by loss of GCase activity may also contribute to this chain of toxic events triggered by *GBA* mutations and involving αS. This possibility is supported by findings showing that αS-lipid interactions modulate protein aggregation, can promote αS assembly and accelerate the rate of αS amyloid formation (Zhu and Fink, 2003; Martinez *et al*., 2007; Franceschi *et al*., 2011; Galvagnion *et al*., 2015, 2016; Grey *et al*., 2015; Gaspar *et al*., 2018).

The aim of the present study was twofold. First, we performed a comprehensive lipidomic analysis of membrane preparations from human fibroblasts and compared data in cells from control individuals *versus* cells from iPD patients without *GBA* mutations and PD-GBA patients carrying the L444P mutation. Results revealed that loss of GCase activity in PD-GBA fibroblasts was associated with a specific lipid profile. This profile featured differences in SL levels and SL acyl chain composition, including a higher proportion of molecules with shorter hydrocarbon chain length. Subsequent experiments assessed the potential link between PD-GBA-related lipid changes and αS aggregation. Findings provided evidence of such a relationship, showing accelerated formation of amyloid fibrils after incubations of αS in the presence of lipids extracted from PD-GBA fibroblasts. Data also showed that co-assembly of SL with αS occurred during the fibrillation process and involved preferentially SL with shorter chain length.

## MATERIALS AND METHODS

### Fibroblasts

Collection and use of human tissue were done in agreement with the principles of the Declaration of Helsinki. Fibroblasts were obtained from skin biopsies that were performed at the IRCCS Mondino Foundation under a research protocol previously approved by the institutional Ethic Committee. Informed consent was obtained from all subjects who underwent the procedure. Fibroblast cultures were grown in RPMI-1640 Medium (Sigma Aldrich) with 10% serum (Sigma Aldrich), 2 mM L-Glutamine (Sigma Aldrich), 100 µg/mL streptomycin and 100 units/mL penicillin. Analyses were carried out at low culture passages, and disease and control cultures were matched for passage number.

### GCase activity assay

Fibroblasts were trypsinized and re-suspended in lysis buffer (0.1 M sodium citrate, 0.1% Triton X-100, 6.7 mM sodium taurocholate). The resulting cell suspensions were lysed by sonication at 4°C, and lysates were incubated on ice for 20 minutes before being centrifuged at 15,000 g for 15 minutes. The supernatants were collected into new tubes and their protein content determined using bicinchoninic acid assay. Lysates were then diluted to a protein concentration of 0.1 µg/µL with assay buffer (0.1 M sodium citrate, 0.1% Triton X-100, 6.5 mM sodium taurocholate, 2.5 mM 4-methylumbelliferyl β-D-glucopyranoside) and incubated in non-binding plates (Corning 3881) while shaking (300 rpm) at 37°C. Fluorescence was measured using a FLUOstar Omega plate reader (BMG) with excitation/emission filters of 355-20 / 460 nm. The fluorescence of 25 µM 4-methylumbelliferone was measured under the same conditions and used to convert fluorescence units into µM of 4-methylumbelliferone. GCase activity was calculated using the following equation:

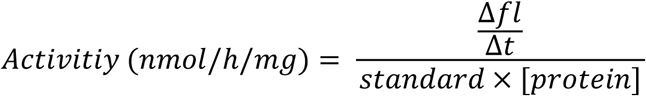

Where 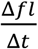 is the change in fluorescence intensity per hour (fluorescence unit / h), standard is the fluorescence (fluorescence unit / µM) of 4-methylumbelliferone and [protein] is the total protein concentration in the sample (0.1 mg/mL).

### Western Blot analysis

Fibroblasts were trypsinized and resuspended in lysis buffer with protease inhibitors. Cell lysates were incubated on ice with shaking and then centrifuged at 16,000 g for 15 min. The supernatants were transferred to fresh tubes, and lysates containing 30 µg of protein were electrophoresed on a NuPageTM 4–12% Bis-Tris Protein gel (Thermo Fisher Scientific). Proteins were transferred to a PVDF membrane (Amersham), blocked in 5% BSA and treated with primary and secondary antibodies. Antibody binding was detected using an ECL chemiluminescence kit. The following antibodies were used: glucocerebrosidase (ab55080 Abcam, dilution 1:1000) and β-actin (ab8227, Abcam, dilution 1:7500).

### Lipid extraction for mass spectrometry lipidomics

Mass spectrometry-based lipid analysis was performed by Lipotype GmbH (Dresden, Germany) as described (Sampaio *et al*., 2011). Lipids were extracted using a two-step chloroform/methanol procedure (Ejsing *et al*., 2009). Samples were spiked with internal lipid standard mixture containing: cardiolipin 16:1/15:0/15:0/15:0 (CL), Cer 18:1;2/17:0, diacylglycerol 17:0/17:0 (DAG), hexosylceramide 18:1;2/12:0 (HexCer), lyso-phosphatidate 17:0 (LPA), lyso-phosphatidylcholine 12:0 (LPC), lyso-phosphatidylethanolamine 17:1 (LPE), lyso-phosphatidylglycerol 17:1 (LPG), lyso-phosphatidylinositol 17:1 (LPI), lyso-phosphatidylserine 17:1 (LPS), phosphatidate 17:0/17:0 (PA), phosphatidylcholine 17:0/17:0 (PC), phosphatidylethanolamine 17:0/17:0 (PE), phosphatidylglycerol 17:0/17:0 (PG), phosphatidylinositol 16:0/16:0 (PI), phosphatidylserine 17:0/17:0 (PS), cholesterol ester 20:0 (CE), sphingomyelin 18:1;2/12:0;0 (SM), triacylglycerol 17:0/17:0/17:0 (TAG) and cholesterol D6 (Chol). After extraction, the organic phase was transferred to an infusion plate and dried in a speed vacuum concentrator. First-step dry extract was re-suspended in 7.5 mM ammonium acetate in chloroform/methanol/propanol (1:2:4, V:V:V) and 2^nd^-step dry extract in 33% ethanol solution of methylamine in chloroform/methanol (0.003:5:1; V:V:V). All liquid handling steps were performed using Hamilton Robotics STARlet robotic platform with the Anti Droplet Control feature for organic solvents pipetting.

### MS data acquisition

Samples were analysed by direct infusion on a QExactive mass spectrometer (Thermo Scientific) equipped with a TriVersa NanoMate ion source (Advion Biosciences). Samples were analysed in both positive and negative ion modes with a resolution of Rm/z=200=280000 for MS and Rm/z=200=17500 for MSMS experiments, in a single acquisition. MSMS was triggered by an inclusion list encompassing corresponding MS mass ranges scanned in 1 Da increments (Surma *et al*., 2015). Both MS and MSMS data were combined to monitor CE, DAG and TAG ions as ammonium adducts; PC, PC O-, as acetate adducts; and CL, PA, PE, PE O-, PG, PI and PS as deprotonated anions. MS only was used to monitor LPA, LPE, LPE O-, LPI and LPS as deprotonated anions; Cer, HexCer, SM, LPC and LPC O-as acetate adducts and cholesterol as ammonium adduct of an acetylated derivative (Liebisch *et al*., 2006).

### Data analysis and post-processing

Data were analysed with in-house developed lipid identification software based on LipidXplorer (Herzog *et al*., 2011, 2012). Data post-processing and normalization were performed using an in-house developed data management system. Only lipid identifications with a signal-to-noise ratio >5, and a signal intensity 5-fold higher than in corresponding blank samples were considered for further data analysis.

### Aggregation kinetics measurements

Recombinant WT *α*S was produced as previously described (Galvagnion *et al*., 2015). For each fibroblast line, lipids were extracted from one million cells resuspended in 200 μL PBS using 1 mL chloroform:methanol mixture (10:1, v:v). The organic phase was then evaporated and the lipids incubated in the presence of 50 μM *α*S and 50 μM Thioflavin-T in MES buffer (10 mM MES, pH 5.0, 0.01% sodium azide) at 37°C in high binding plates (Corning 3601) under quiescent conditions. The fluorescence intensity was measured on a FLUOstar Omega plate reader (BMG) with excitation/emission filters of 448-10 / 482-10 nm.

### Statistics

Data are expressed as mean ± SEM (standard error of the mean). Unpaired t-test (two-tailed P value) was used for comparisons of means between two groups (control *vs*. PD, PD *vs*. PD-GBA and control *vs*. PD-GBA). Statistical analysis was performed using GraphPad Prism v8, GraphPad Software, CA, USA. Correlations were analysed with Pearson correlation analysis. Statistical significance was set at P < 0.05.

## RESULTS

### *GBA* mutation and GCase activity in human fibroblasts

Fibroblasts were generated from skin biopsies of 4 control subjects with no history of neurological disorders (2 males and 2 females), 4 iPD patients (2 males and 2 females) and 5 PD-GBA patients carrying a heterozygous L444P *GBA* mutation (3 males and 2 females). Average age of onset of clinical manifestations was 57 ± 3.5 and 47 ± 4.3 years in the iPD and PD-GBA groups, respectively, consistent with an earlier disease onset of GBA-associated PD (Schapira, 2015). GCase protein levels, as assessed by western blot analysis of lysed cells, were unchanged between the control and the two patient groups (Fig. 2A and B). GCase enzyme activity was also similar in fibroblasts from control subjects and iPD patients, but was significantly decreased by approximately 25% in fibroblasts from PD-GBA patients (Fig. 2C).

**Figure 2.**
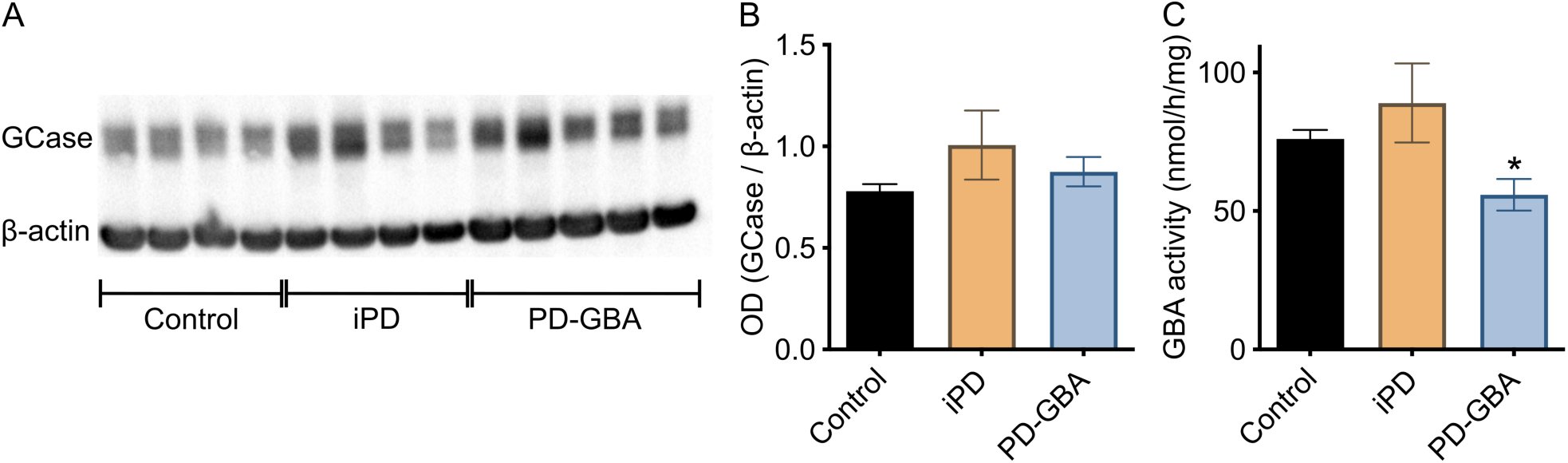
GCase protein levels and activity in fibroblasts. Measurements were made in fibroblasts from control subjects and iPD and PD-GBA patients. (**A**) Western blots showing immunoreactivity for human GCase and β-actin. (**B**) Semi-quantitative analysis of band intensities of the blots shown in panel A. (**C**) Measurements of GCase activity in fibroblasts from the control (*n* = 4), iPD (*n* = 4) and PD-GBA (*n* = 5) groups. Bars show mean values, and error bars are ± SEM. Unpaired *t* test comparing data in control subjects *vs*. values in the iPD or PD-GBA groups. Control *vs*. PD-GBA (*): *P* = 0.0258; F(4,3) = 3.734.

### Lipidome of human fibroblasts

Mass spectrometry after direct infusion (“shotgun lipidomics”) was used for the first time to perform a comprehensive quantitative and qualitative analysis of the lipid composition of fibroblasts from control subjects and Parkinson’s Disease patients with and without a *GBA* mutation. The lipidome of control cells mainly consisted of phospholipids (74.5% of total lipids), Chol (19%), SL (4.2%) and glycerides (1.6%) (Fig. 3A-D). A detailed quantification of phospholipid species is reported in Supplementary Fig. 1. The 4.2 per cent of total SL was comprised of 3.6%, 0.4% and 0.2% SM, Cer and HexCer, respectively (Fig 3E-G).

**Figure 3.**
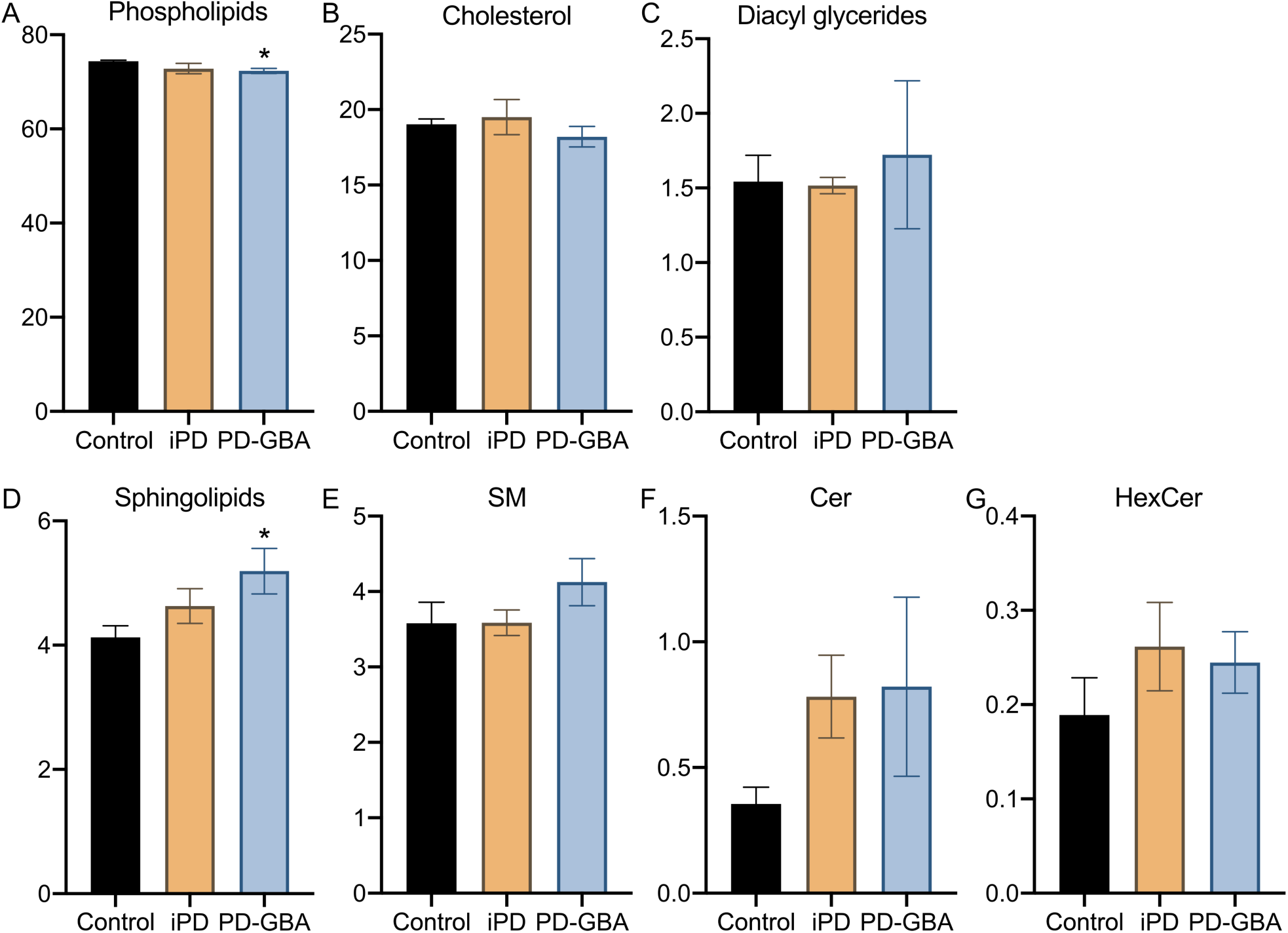
Lipid levels in control, iPD and PD-GBA fibroblasts. Levels of (**A**) phospholipids, (**B**) cholesterol, (**C**) diacyl glycerides and (**D**) total and (**E-G**) specific ((**E**) sphingomyelin, SM, (**F**) ceramide, Cer and (**G**) hexosylceramide, HexCer) sphingolipids were assayed in fibroblast lipid extracts. Measurements were made in extracts from control subjects (*n* = 4) and iPD (*n* = 4) and PD-GBA (*n* = 5) patients. Data are expressed as percentage of the total lipid content. Bars show mean values, and error bars are ± SEM. Unpaired *t* test was performed to compare data in the control *vs*. iPD and control *vs*. PD-GBA groups (*). Control *vs*. PD-GBA (*): Phospholipids [*P* = 0.0130; F(4,3) = 7.330] and sphingolipids [*P* = 0.0485; F(4,3) = 4.766].

Fibroblast lipid composition did not differ significantly between control subjects and iPD patients (Fig. 3 and Supplementary Fig. 1). Specific changes were instead observed in the lipidome of cells from PD-GBA patients. In these cells, the most prominent variation concerned the levels of total SL that were significantly increased by approximately 25% (Fig. 3D). The percent of specific SL species, namely SM, Cer and HexCer, also tended to be higher in PD-GBA fibroblasts, although these changes did not reach statistical significance (Fig. 3E-G). Levels of cholesterol and glycerides were similar to control values, whereas a small but significant 3% decrease in the percent of total phospholipids was detected in samples from PD-GBA patients (Fig. 3A).

### Chemical properties of sphingolipids in human fibroblasts

Our lipidomic analysis of human fibroblasts also assessed levels of specific SL molecules and, in particular, identified and quantified SM, Cer and HexCer molecules with different hydrocarbon chain length and degree of unsaturation. Data showed that most SL (80-90% of SM, Cer and HexCer) had either 34 (C34) or 42 (C42) hydrocarbons (Fig. 4A, C and D). SM with 34 hydrocarbons and a single double bond (SM 34:1; 18:1/16:0) and SM with 42 hydrocarbons and two double bonds (SM 42:2; 18:1/24:1) were the two main SM species detected in our fibroblast preparations; they accounted for approximately 55% and 15%, respectively, of the total SM levels (Fig. 4B). Other abundant SL molecules were Cer 34:1 (18:1/16:0) and Cer 42:1 (18:1/24:0), which accounted for 40% and 25% of all Cer (Fig. 4D), and HexCer 34:1 (18:1/16:0) and HexCer 42:1 (18:1/24:0), representing 20% and 35% of the total HexCer (Fig. 4F).

**Figure 4.**
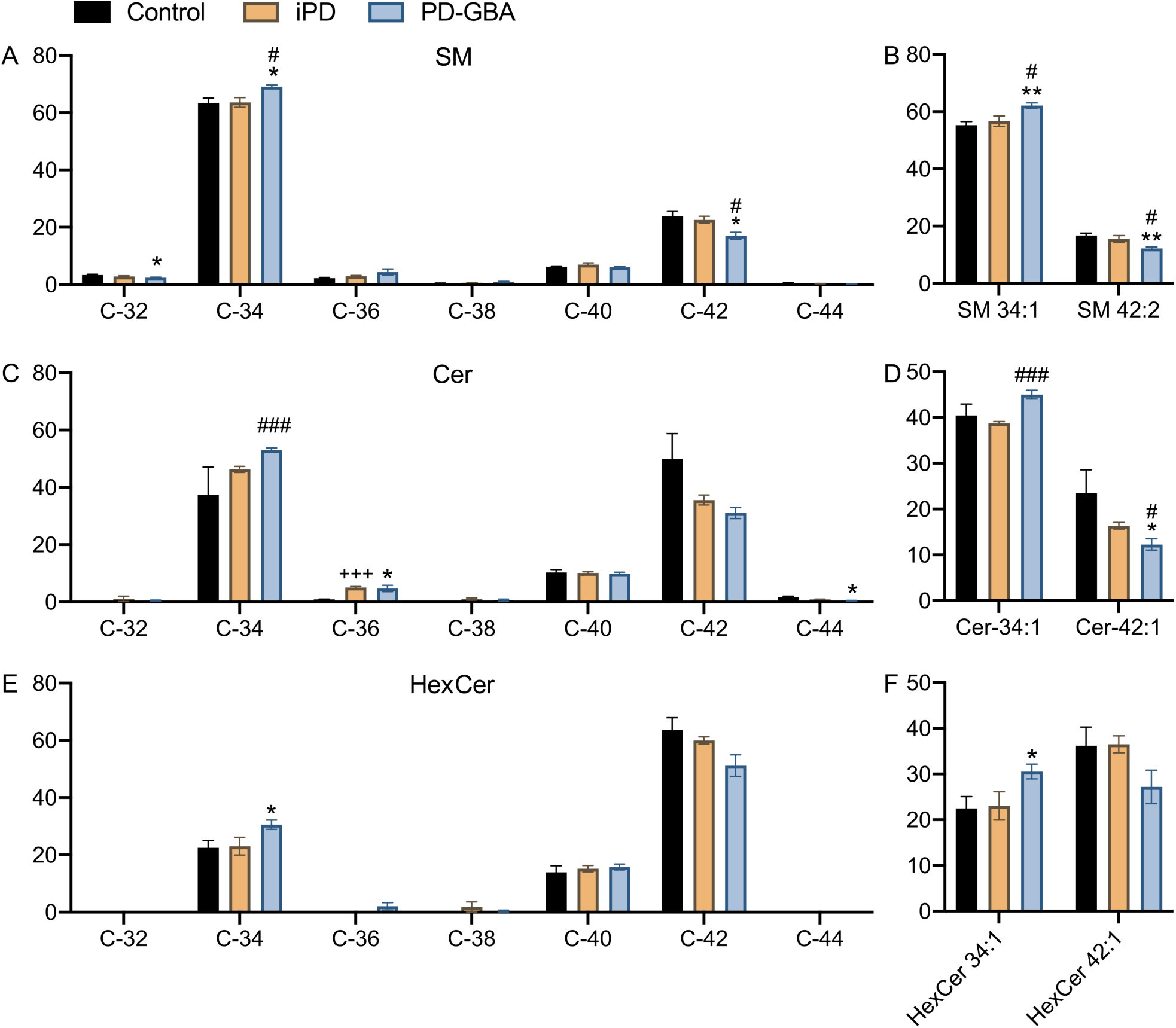
Levels of sphingolipid molecules with different acyl chain length. Measurements of sphingomyelin (SM), ceramide (Cer) and hexosylceramide (HexCer) molecules were made in fibroblast lipid extracts from control subjects (*n* = 4) and iPD (*n* = 4) and PD-GBA (*n* = 5) patients. (**A, C** and **E**) Levels of SM, Cer and HexCer with different hydrocarbon chain lengths (C-32 to C-44) are shown as percent of the respective SM, Cer and HexCer total content. Bars show mean values, and error bars are ± SEM. Multiple *t* test was performed to compare means between two groups, control *vs*. iPD (^+^), control *vs*. PD-GBA (*) and iPD *vs*. PD-GBA (^#^). *^,#^*P* < 0.05; ^+++, ###^*P* < 0.001. Control *vs*. PD-GBA (*): SM C-32 (*P* = 0.0269), SM C-34 (*P* = 0.0113), SM C-42 (*P* = 0.0154), Cer C-36 (*P* = 0.0188), Cer C-44 (*P* = 0.0197), HexCer C-34 (*P* = 0.0285). iPD *vs*. PD-GBA (^#, ###^): SM C-34 (*P* = 0.0124), SM C-42 (*P* = 0.0144), Cer C-34 (*P* < 0.001). Control *vs*. iPD (^+++^): Cer C-36 (*P* < 0.001). (**B**) Levels of the two main SM molecules, SM 34:1 and SM 42:2, are shown as percent of the total SM content. ^#^*P* < 0.05; \*\**P* < 0.005. Control *vs*. PD-GBA (**): SM 34:1 (*P* = 0.0034), SM 42:2 (*P* = 0.0024). iPD *vs*. PD-GBA (^#^): SM 34:1 (*P* = 0.0258), SM 42:2 (*P* = 0.0282). (**D**) Levels of the two main Cer molecules, Cer 34:1 and Cer 42:1, are shown as percent of the total Cer content. *^,#^*P* < 0.05; ^###^*P* < 0.001. Control *vs*. PD-GBA (*): Cer 42:1 (*P* = 0.0477). PD *vs*. PD-GBA (^#, ###^): Cer 34:1 (*P* < 0.001), Cer 42:1 (*P* = 0.0329). (**F**) Levels of the two main HexCer molecules, HexCer 34:1 and HexCer 42:1, are shown as percent of the total HexCer content. \**P* < 0.05. Control *vs*. PD-GBA (*): HexCer 34:1 (*P* = 0.029).

Potential changes in SL with shorter and longer hydrocarbon chains were then compared between control cells and fibroblasts from iPD and PD-GBA patients. Levels of C34 and C42 sphingolipids were not significantly different in control *vs*. iPD-derived preparations (Fig. 4A, C and E). In contrast, when measurements were compared between PD-GBA *vs*. control or iPD fibroblasts, data revealed an increase in short-chain (C34) SM, Cer and HexCer and a consistent reduction of long-chain (C42) SM, Cer and HexCer (Fig. 4A, C and E). A comparison of levels of specific SL molecules confirmed these differences. SM 34:1, Cer 34:1 and HexCer 34:1 were all increased by 15-35% in mutation-carrying *vs*. control and/or iPD fibroblasts (Fig. 4B, D and F). In the same PD-GBA cells, measurements of SM 42:2, Cer 42:1 and HexCer 42:1 showed a decrease that ranged between 25% and 50% (Fig. 4B, D and F).

### Fibroblast sphingolipid composition and GCase activity

Taken together, results obtained from our lipidomic analysis indicated that loss of GCase activity in PD-GBA fibroblasts was accompanied not only by an overall increase in SL levels but also higher levels of short-chain and lower levels of long-chain SL. In these cells, the ratios C34:C42 SM, C34:C42 Cer and C34:C42 HexCer were indeed increased by 50%, 95% and 70%, respectively (Fig. 5A-C). To assess whether changes in acyl chain length were specific for SL molecules, levels of phospholipids with different carbon numbers were also measured and compared in control subjects and iPD and PD-GBA patients (Supplementary Fig. 2). Data showed small differences in the percent of specific phospholipid molecules among the three groups but no trend toward an increase in short-chain and decrease in long-chain phospholipids in fibroblast extracts from PD-GBA patients (Supplementary Fig. 2). To evaluate further the relationship between SL composition and GCase activity, C34:C42 SM, Cer and HexCer ratio values were calculated for each control, iPD and PD-GBA cell preparation and then plotted against the corresponding values of enzyme activity. This analysis revealed a significant inverse correlation. Higher enzyme activity was associated with a lower ratio and *vice versa*, supporting the conclusion that changes in GCase activity alter the content and relative proportion of short- and long-chain SL (Fig. 5D-F). The increase in C34:C42 SL ratio, as detected in our study, is therefore likely to be a specific consequence of reduced GCase activity and a feature of altered lipid metabolism caused by the L444P *GBA* mutation.

**Figure 5.**
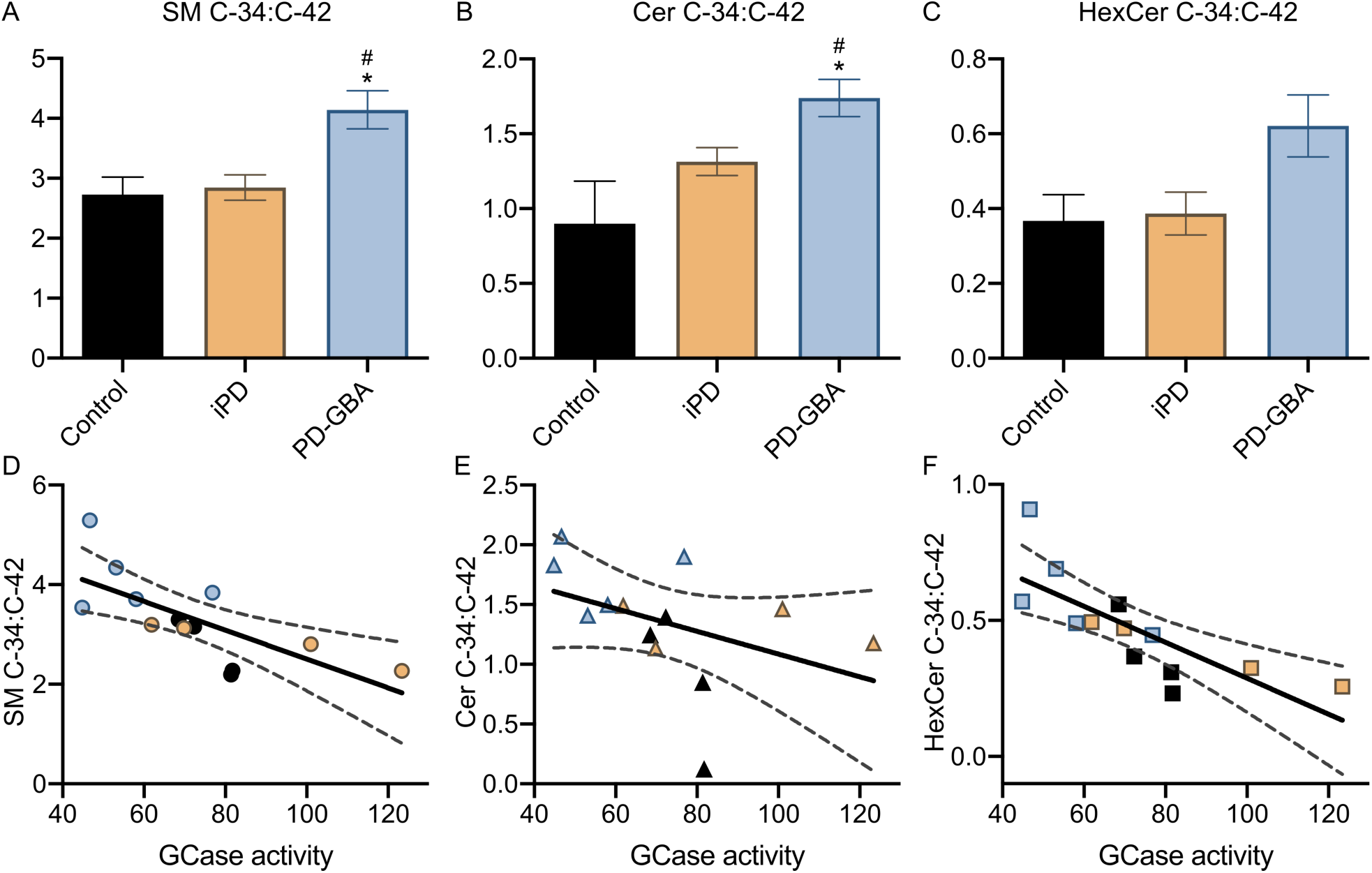
Correlation between the ratio short-over long-chain sphingolipids and GCase activity. (**A**-**C**) The ratios C-34:C-42 (percent C-34 over percent C-42) SM, C-34:C-42 Cer and C-34:C-42 HexCer were calculated from lipid measurements made in fibroblasts from control subjects (*n* = 4) and iPD (*n* = 4) and PD-GBA (*n* = 5) patients. Bars show mean values, and error bars are ± SEM. Unpaired *t* test was performed to compare data in control *vs*. PD-GBA (*) and iPD *vs*. PD-GBA (^#^). *^,#^*P* < 0.05. Control *vs*. PD-GBA (*): SM C-34:C:42 [P = 0.0147; F(4,3) = 1.504], Cer C34:C-42 [*P* = 0.0221, F(3,4) = 4.173]. iPD *vs*. PD-GBA (^#^): SM C-34:C:42 [*P* = 0.0149; F(4,3) = 2.791], Cer C34:C-42 [*P* = 0.0352, F(4,3) = 2.246]. (**D**-**F**) C34:C42 SM, Cer and HexCer ratio values for each control (*n* = 4, black), iPD (*n* = 4, orange) and PD-GBA (*n* = 5, blue-gray) fibroblast preparation were plotted against the corresponding values of GCase activity. Pearson correlation analysis was performed to assess the strength of the association between: C-34:C:42 SM ratio and GCase activity (*P* = 0.0053; r = −0.7226), C-34:C-42 Cer ratio and GCase activity (*P* = 0.1546; r = −0.4186) and C-34:C-42 HexCer ratio and GCase activity (*P* = 0.0020; r = −0.7729).

### Relationship between fibroblast lipid composition and α-synuclein aggregation

Specific lipid-αS interactions modulate αS aggregation (Zhu and Fink, 2003; Martinez *et al*., 2007; Franceschi *et al*., 2011; Galvagnion *et al*., 2015, 2016; Grey *et al*., 2015; Gaspar *et al*., 2018). The next set of experiments was therefore designed to test the hypothesis that differences in lipid composition observed in fibroblasts from PD-GBA patients may affect αS *in vitro* fibrillation. For these experiments, lipids were obtained from control (LIPID_control_) and PD-GBA (LIPID_PD-GBA_) fibroblasts using the same extraction procedure that was employed for our lipidomic analysis. The kinetics of αS aggregation was monitored over a 90-min incubation time following the shift in fluorescence caused by the binding of thioflavin T (ThT) to amyloid structures. Addition of fibroblast-derived lipids, which were resuspended as vesicles, triggered αS aggregation as indicated by an increase in ThT fluorescence. Interestingly, kinetics curves of the ThT signal generated from PD-GBA samples were distinctly shifted to the left, with fluorescence rising and reaching its plateau at earlier time points (Fig. 6A). When reaction half-times (t ½) were plotted and compared between incubations of αS with LIPID_control_ *vs*. LIPID_PD-GBA_, a significantly lower t ½ was observed under the latter condition, consistent with a pro-aggregation effect of LIPID_PD-GBA_ (Fig. 6B).

**Figure 6.**
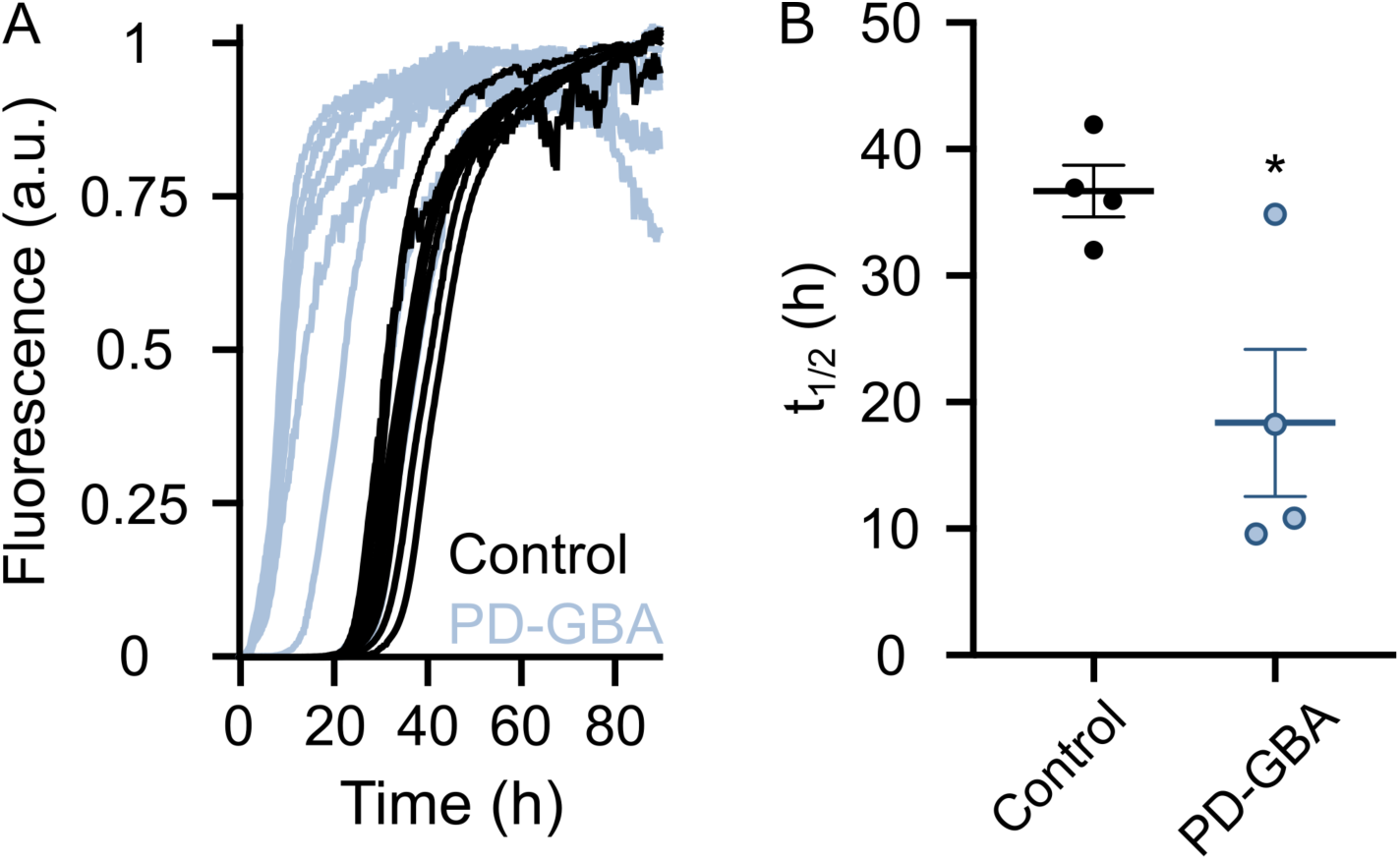
Kinetics of α-synuclein aggregation in the presence of fibroblast lipid extracts. (**A**) Thioflavin T fluorescence was measured as an indicator of amyloid fibril formation during incubations of α-synuclein in the presence of lipid extractrs from control (*n* = 4; black) and PD-GBA (*n* = 4; blue-gray) fibroblast preparations. Kinetics curves are shown from two separate incubations of each subject/patient extract. (**B**) Half-times of the reaction of α-synuclein fibril formation after addition of lipid extracts from control subjects (*n* = 4) and PD-GBA patients (*n* = 4). Each circle represents the average of duplicate measurements. Bars show mean values, and error bars are ± SEM. Unpaired *t* test was performed to compare data. \**P* = 0.0248; F(3,3) = 8.102.

Amyloid fibrils that were formed after incubations of αS with LIPID_control_ or LIPID_PD-GBA_ were collected (at the time when ThT signals reached their plateau) and subjected to lipid extraction. Solvent mixtures were then analysed using mass spectrometry to determine if specific lipid molecules were co-assembled with αS fibrils (Hellstrand *et al*., 2013; Galvagnion *et al*., 2019).

Results showed that the composition of fibril-derived lipid mixtures consisted mainly of PC (approximately 75% of total lipids), DAG (10%) and SL (12%); the 12 per cent of total SL was comprised of 7.1%, 1.4% and 3.4% SM, Cer and HexCer, respectively (Fig. 7A). Lipid composition of αS fibrils did not differ significantly between samples collected after incubations with LIPID_control_ or LIPID_PD-GBA_ (Fig. 7A). Fibril lipid mixtures were also analysed for the presence of short- and long-chain SL. Similar to results obtained from the lipidomic analysis of whole fibroblasts, most SL extracted from αS fibrils had 34 or 42 hydrocarbons (Fig. 7B-D). The relative proportion of short (C34)- and long (C42)-chain molecules was different, however, between whole fibroblast-*vs*. fibril-derived samples (cf. data in Fig. 4 and Fig. 7B-D). In particular, marked changes were observed in the percent of short- and long-chain SM and HexCer. C34 SM and C34 HexCer represented approximately 60% and 20% of the total SM and HexCer levels in extracts of whole fibroblasts (Fig. 4A, G); they instead accounted for >90% of SM and HexCer that were extracted from amyloid fibrils after incubations of αS with either LIPID_control_ or LIPID_PD-GBA_ (Fig. 7B, D). On the other hand, levels of C42 SM and C42 HexCer, which accounted for a significant percent of total SM and HexCer in whole fibroblasts, were barely detected in lipid mixtures from αS fibrils. Taken together, these data indicate an enrichment of C34 SM and HexCer in fibril-derived lipid extracts and are consistent with a high propensity of these short-chain molecules to interact with and be incorporated into amyloid fibrils during the process of αS aggregation.

**Figure 7.**
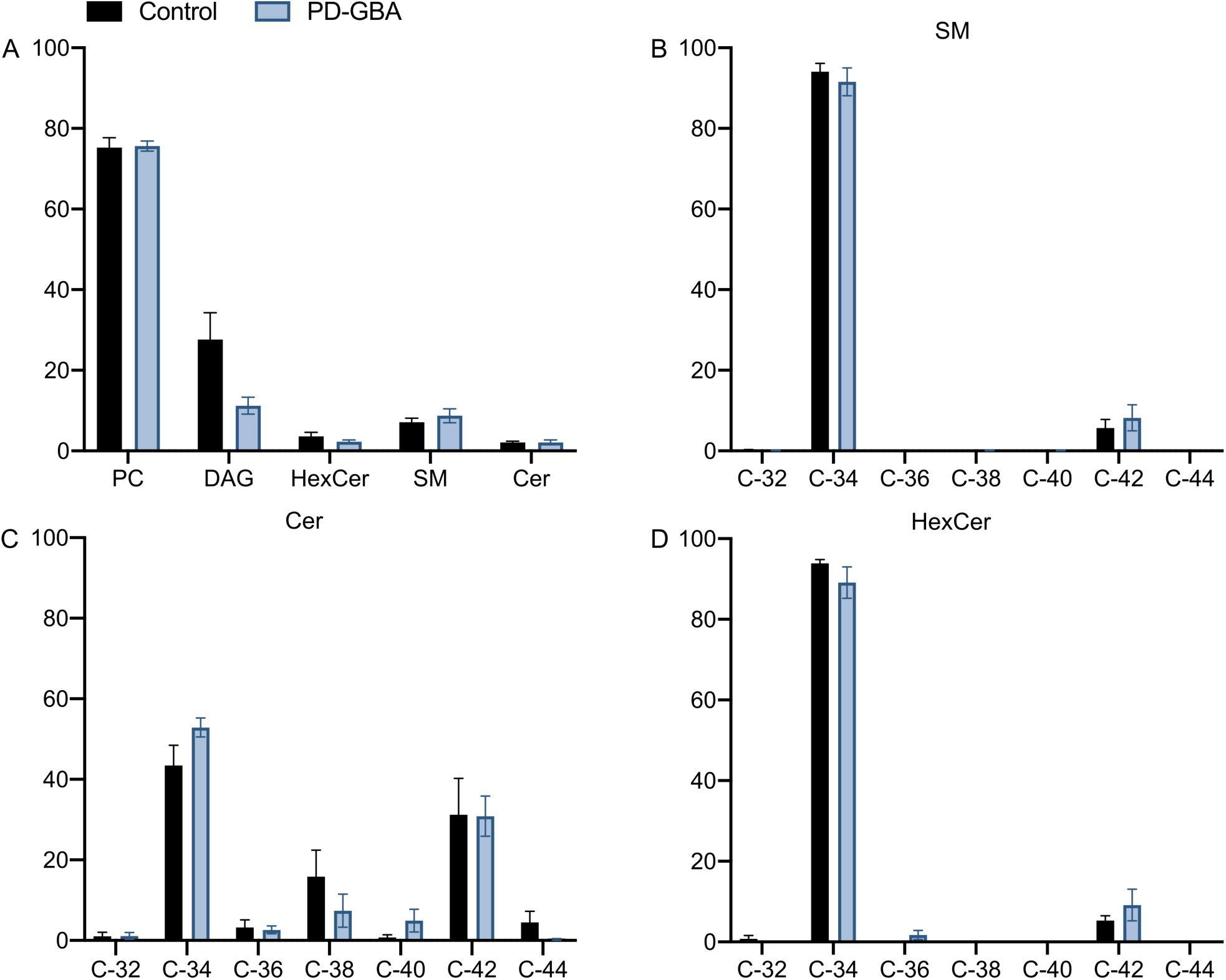
Lipid composition of α-synuclein fibrils. Amyloid fibrils were collected at the end of incubations of α-synuclein in the presence of fibroblast lipid extracts from control subjects (*n* = 4) and PD-GBA patients (*n* = 4). These fibrils were subjected to lipid extraction, and fibril lipid content was measured by mass spectrometry. (**A**) Data show the percent of lipid species, phosphatidylcholine (PC), diacyl glyceride (DAG), hexosylceramide (HexCer), sphingomyelin (SM) and ceramide (Cer), detected in extracts from α-synuclein fibrils. (**B**-**D**) Levels of SM, Cer and HexCer with different hydrocarbon chain lengths (C-32 to C-44) are shown as percent of the respective SM, Cer and HexCer total content. Bars show mean values, and error bars are ± SEM.

## DISCUSSION

Results of this study indicate the potential value of patient-derived fibroblasts as an *ex vivo* system for the identification and monitoring of changes in lipid metabolism caused by *GBA* mutations. A unique lipid profile was detected by shotgun lipidomics of cultures from carriers of the severe L444P *GBA* mutation; this profile distinguished PD-GBA fibroblasts from cells that were obtained from control subjects as well as cells from iPD patients without *GBA* mutations. Detailed analyses revealed not only that levels of SL were increased but also that the ratio of short-over long-chain SL was distinctly altered in fibroblasts from mutation carriers, thus providing a new biological fingerprint of the mutation in these Parkinson’s Disease patients. An additional important finding of this study was that membrane lipids isolated from PD-GBA cultures were significantly more effective than lipids from control cells in triggering *in vitro* fibrillation of recombinant human αS. This finding supports a mechanism of likely pathophysiological relevance linking changes in cell lipid composition caused by *GBA* mutations to αS aggregate pathology.

A correlation was found between the unique lipid profile of PD-GBA fibroblasts and the activity of GCase that was significantly decreased in L444P mutation carriers as compared to control subjects and iPD patients. This latter finding, i.e. lack of GCase changes in Parkinson’s Disease patients without a *GBA* mutation, is consistent with data of earlier studies using control- and patient-derived fibroblasts (McNeill *et al*., 2014; Ambrosi *et al*., 2015; Sanchez-Martinez *et al*., 2016; Collins *et al*., 2018). It is apparently at odds, however, with the results of previous investigations in which GCase activity was measured and found to be lowered in brain tissue specimens from iPD as compared to control subjects (Gegg *et al*., 2012; Rocha *et al*., 2015; Huebecker *et al*., 2019). In contrast to neurons, fibroblasts do not express αS. Thus, a plausible explanation for the different results in fibroblasts and brain tissue may be that, in the latter, loss of GCase activity specifically arises from a pathological accumulation of αS (Schapira *et al*., 2014). Consistent with this interpretation, a region-specific reduction of GCase activity has been reported in the brain of patients with sporadic PD; this effect was correlated with αS burden and early development of αS pathology (Murphy and Halliday, 2014).

Reduced GCase activity in PD-GBA fibroblasts significantly affected SL content and metabolism, resulting in higher levels of total SL, enhanced amounts of short-chain SL molecules and decreased content of long-chain SL. Earlier studies have reported accumulation of SL and, in particular, GluCer and GluSph as a consequence of reduced GCase activity in cell lines (i.e. iPSC-derived neurons) from Gaucher’s Disease and PD-GBA patients (Schöndorf *et al*., 2014; Aflaki *et al*., 2016). Our current data not only extend these observations but reveal for the first time that changes in acyl chain length of SL significantly contribute to the altered lipid profile of PD patients with *GBA* mutations. Interestingly, the hydrocarbon chain length of Cer, HexCer and SM molecules were all affected by a loss of GCase activity, underscoring the interrelated nature of SL metabolism and suggesting that an initial GCase impairment ultimately alters the function of other enzymes involved in this metabolism. Indeed, as indicated by the results of previous investigations, a shift from long- to short-chain SL may result from changes in the expression/activity of enzymes such as α-sphingomyelinase and ceramide synthases (CerS) (Imgrund *et al*., 2009; Ben-David *et al*., 2011; Mosbech *et al*., 2014; van Smeden *et al*., 2014). For example, accumulation of short-chain C18 GluCer was observed in different brain regions of a Gaucher’s Disease mouse model (4L/PS/NA) homozygous for a mutant GCase (V394L [4L]) and expressing a prosaposin hypomorphic (PS-NA) transgene; this accumulation appeared to be correlated with the regional distribution of CerS1 and CerS2 (Jones *et al*., 2017). It is also noteworthy that a significant shift in Cer acyl chain composition toward shorter chain length was measured in the anterior cingulate cortex of brains from iPD patients and attributed to an upregulation of CerS1 expression (Abbott *et al*., 2014).

SL are bioactive molecules playing both structural and signalling roles in a range of cellular processes including cell growth, differentiation and migration, autophagy, inflammation, response to trophic factors and apoptosis (Lingwood and Simons, 2010; Lippincott-Schwartz and Phair, 2010; Hannun and Obeid, 2018). Thus, by altering cellular SL composition, *GBA* mutations could have broad functional and pathological consequences, most of all within the brain where SM, HexCer and Cer molecules are particularly abundant and tightly regulated (O’Brien and Sampson, 1965; Olsen and Færgeman, 2017). Structural and signalling functions of SL are also significantly affected by their saturation and length of acyl chain. In particular, accumulation of short-chain at the expenses of long-chain SL could alter the structural order of membrane lipid bilayers, modify membrane curvature and disrupt membrane fluidity (Niemelä *et al*., 2006; Ben-David and Futerman, 2010; Mencarelli and Martinez-Martinez, 2013). A switch from long- to short-chain molecules could also interfere with lipid-protein interactions at the plasma membrane level and affect signalling responses involved, for example, in lipid and protein trafficking and degradation and in cell death pathways (Sillence *et al*., 2002; Kroesen *et al*., 2003; Koivusalo *et al*., 2007; Sassa *et al*., 2012; Ali *et al*., 2013; Backman *et al*., 2018).

Implications of the results of this study on mechanisms of αS pathology are particularly relevant not only due to the important role played by αS in Parkinson’s Disease pathogenesis but also in view of evidence linking *GBA* mutations to αS alterations. Findings of earlier investigations strongly support a relationship between GCase deficiency, increased levels of SL and accumulation and aggregation of αS in cells treated with a GCase inhibitor, iPSC-derived DA neurons from Gaucher’s Disease and PD-GBA patients and brain tissue from Gaucher’s Disease mouse models and *GBA*-mutant mice (Sun *et al*., 2005; Mazzulli *et al*., 2011; Sardi *et al*., 2011; Xu *et al*., 2011; Cleeter *et al*., 2013; Schöndorf *et al*., 2014; Aflaki *et al*., 2016; Migdalska-Richards *et al*., 2017). An important role of lipid-αS interactions in pathogenetic processes triggered by *GBA* mutations is also suggested by earlier biochemical and biophysical studies showing that the chemical properties of lipids significantly affect the binding of αS to membranes and the kinetics of membrane-induced αS aggregation (Galvagnion, 2017). αS binds preferentially to membranes in a fluid state and with high curvature (Middleton and Rhoades, 2010; Galvagnion *et al*., 2016). Therefore, changes in membrane lipid composition that, as discussed above, could alter these biophysical features could also interfere with αS function and promote its toxic potential.

Synthetic membrane preparations have been effectively used to gain insight into the properties of membranes with different lipid compositions. In an earlier study, formation of αS amyloid fibrils was compared in the presence of synthetic membranes composed of lipids with the same head group, i.e. phosphatidylserine, but with hydrocarbon chains of different length. The reported results showed that αS aggregation was significantly enhanced in the presence of lipids with shorter hydrocarbon chains (Galvagnion *et al*., 2016). Our present findings are in line with these earlier observations. Fibril formation was indeed accelerated when αS was incubated with membrane lipids from PD-GBA cells that contained higher levels of Cer, HexCer and SM molecules with shorter hydrocarbon chains. Short chain SM and HexCer species were also found to be enriched in lipid extracts from αS fibrils collected at the end of lipid/αS incubations. This latter new finding suggests that short chain SL have a high propensity to co-assemble with αS fibrils and may act as preferred reactants during lipid-induced αS amyloid formation.

Extrapolation of results from *ex vivo* models (patient-derived fibroblasts) and *in vitro* experiments (incubations of αS with membrane lipids) to pathological processes always requires caution. Nevertheless, taken together, data of our study support the concept that changes in lipid chemical properties induced by *GBA* mutations are of likely pathophysiological relevance. Data suggest, for example, that an altered lipid profile induced by loss of GCase activity may itself represent a risk factor for the development of αS aggregate pathology in carriers of *GBA* mutations. Findings of this study and follow-up investigations into the effects of *GBA* mutations on fibroblast lipid content also bear potential clinical implications. SL changes could conceivably be more pronounced in fibroblasts from PD-GBA patients as compared to aged-matched carriers without clinical manifestations. They may therefore be considered for diagnostic purposed as a biomarker of disease risk and disease conversion. Clinical trials are currently assessing safety and effectiveness of drugs capable of reversing a loss of GCase activity in PD-GBA patients (Mullin *et al*., 2020). As part of these trials, measurements of fibroblast lipid profile could be used as an indicator of therapeutic intervention and response and may help elucidate the relationship between GCase activity, membrane lipid composition and Parkinson’s disease progression.

## ACKNOWLDEGEMENTS

We thank Drs. Roberta Zangaglia and Micol Avenali (IRCCS Mondino Foundation) for their assistance with the enrollment and skin biopsy procedures and Ms. Laura Demmer for her technical assistance with fibroblast cultures and lipid extract preparations.

## SUPPLEMENTARY MATERIAL

See attachment.

## ABBREVIATIONS

αS: α-synuclein
CE: cholesterol ester
Cer: ceramide
CerS: ceramide synthase
Chol: cholesterol
CL: cardiolipin
DAG: diacylglycerol
GBA: glucocerebrosidase 1 gene
GCase: glucocerebrosidase
GluCer: Glucosylceramide
GluSph: glucosylsphingosine
HexCer: hexosylceramide
iPSC: induced pluripotent stem cells
LPA: lyso-phosphatidate
LPC: lyso-phosphatidylcholine
LPE: lyso-phosphatidylethanolamine
LPG: lyso-phosphatidylglycerol
LPI: lyso-phosphatidylinositol
LPS: lyso-phosphatidylserine
PA: phosphatidate
PC: phosphatidylcholine
PE: phosphatidylethanolamine
PG: phosphatidylglycerol
PI: phosphatidylinositol
PS: phosphatidylserine
SL: sphingolipid
SM: sphingomyelin
TAG: triacylglycerol
ThT: thioflavin T

## REFERENCES

Abbott SK, Li H, Muñoz SS, Knoch B, Batterham M, Murphy KE, et al. Altered ceramide acyl chain length and ceramide synthase gene expression in Parkinson’s disease. Mov Disord 2014; 29: 518–26.

Aflaki E, Borger DK, Moaven N, Stubblefield BK, Rogers SA, Patnaik S, et al. A New Glucocerebrosidase Chaperone Reduces -Synuclein and Glycolipid Levels in iPSC-Derived Dopaminergic Neurons from Patients with Gaucher Disease and Parkinsonism. Journal of Neuroscience 2016; 36: 7441–52.

Alcalay RN, Caccappolo E, Mejia-Santana H, Tang M-X, Rosado L, Orbe Reilly M, et al. Cognitive performance of GBA mutation carriers with early-onset PD. Neurology 2012; 78: 1434–40.

Alcalay RN, Levy OA, Waters CH, Fahn S, Ford B, Kuo S-H, et al. Glucocerebrosidase activity in Parkinson’s disease with and without GBA mutations. Brain 2015; 138: 2648–58.

Ali M, Fritsch J, Zigdon H, Pewzner-Jung Y, Schütze S, Futerman AH. Altering the sphingolipid acyl chain composition prevents LPS/GLN-mediated hepatic failure in mice by disrupting TNFR1 internalization. Cell Death Dis 2013; 4: e929.

Ambrosi G, Ghezzi C, Zangaglia R, Levandis G, Pacchetti C, Blandini F. Ambroxol-induced rescue of defective glucocerebrosidase is associated with increased LIMP-2 and saposin C levels in GBA mutant Parkinson’s disease cells. Neurobiol Dis 2015; 82: 235–42.

Anheim M, Elbaz A, Lesage S, Durr A, Condroyer C, Viallet F, et al. Penetrance of Parkinson disease in glucocerebrosidase gene mutation carriers. Neurology 2012; 78: 417–20.

Backman APE, Halin J, Nurmi H, Möuts A, Kjellberg MA, Mattjus P. Glucosylceramide acyl chain length is sensed by the glycolipid transfer protein. PLoS ONE 2018; 13: e0209230.

Bandres-Ciga S, Diez-Fairen M, Kim JJ, Singleton AB. Genetics of Parkinson’s disease: An introspection of its journey towards precision medicine. Neurobiology of Disease 2020; 137: 104782.

Ben-David O, Futerman AH. The role of the ceramide acyl chain length in neurodegeneration: involvement of ceramide synthases. Neuromolecular Med 2010; 12: 341–50.

Ben-David O, Pewzner-Jung Y, Brenner O, Laviad EL, Kogot-Levin A, Weissberg I, et al. Encephalopathy caused by ablation of very long acyl chain ceramide synthesis may be largely due to reduced galactosylceramide levels. J Biol Chem 2011; 286: 30022–33.

Brockmann K, Srulijes K, Hauser AK, Schulte C, Csoti I, Gasser T, et al. GBA-associated PD presents with nonmotor characteristics. Neurology 2011; 77: 276–80.

Cilia R, Tunesi S, Marotta G, Cereda E, Siri C, Tesei S, et al. Survival and dementia in GBA-associated Parkinson’s disease: The mutation matters. Annals of Neurology 2016; 80: 662–73.

Cleeter MW, Chau KY, Gluck C, Mehta A, Hughes DA, Duchen M, et al. Glucocerebrosidase inhibition causes mitochondrial dysfunction and free radical damage. Neurochem Int 2013; 62: 1–7.

Collins LM, Drouin-Ouellet J, Kuan W-L, Cox T, Barker RA. Dermal fibroblasts from patients with Parkinson’s disease have normal GCase activity and autophagy compared to patients with PD and GBA mutations. F1000Res 2018; 6: 1751.

Ejsing CS, Sampaio JL, Surendranath V, Duchoslav E, Ekroos K, Klemm RW, et al. Global analysis of the yeast lipidome by quantitative shotgun mass spectrometry. Proc Natl Acad Sci USA 2009; 106: 2136–41.

Franceschi GD, Frare E, Pivato M, Relini A, Penco A, Greggio E, et al. Structural and Morphological Characterization of Aggregated Species of α-Synuclein Induced by Docosahexaenoic Acid. J Biol Chem 2011; 286: 22262–74.

Futerman AH, Platt FM. The metabolism of glucocerebrosides — From 1965 to the present. Molecular Genetics and Metabolism 2017; 120: 22–6.

Galvagnion C. The Role of Lipids Interacting with alpha-Synuclein in the Pathogenesis of Parkinson’s Disease. J Parkinsons Dis 2017; 7: 433–50.

Galvagnion C, Brown JW, Ouberai MM, Flagmeier P, Vendruscolo M, Buell AK, et al. Chemical properties of lipids strongly affect the kinetics of the membrane-induced aggregation of alpha-synuclein. Proc Natl Acad Sci U S A 2016; 113: 7065–70.

Galvagnion C, Buell AK, Meisl G, Michaels TC, Vendruscolo M, Knowles TP, et al. Lipid vesicles trigger alpha-synuclein aggregation by stimulating primary nucleation. Nat Chem Biol 2015; 11: 229–34.

Galvagnion C, Topgaard D, Makasewicz K, Buell AK, Linse S, Sparr E, et al. Lipid Dynamics and Phase Transition within α-Synuclein Amyloid Fibrils. J Phys Chem Lett 2019: 7872–7.

Gan-Or Z, Amshalom I, Kilarski LL, Bar-Shira A, Gana-Weisz M, Mirelman A, et al. Differential effects of severe vs mild GBA mutations on Parkinson disease. Neurology 2015; 84: 880–7.

Gaspar R, Pallbo J, Weininger U, Linse S, Sparr E. Ganglioside lipids accelerate α-synuclein amyloid formation. Biochimica et Biophysica Acta (BBA) - Proteins and Proteomics 2018; 1866: 1062–72.

Gegg ME, Burke D, Heales SJR, Cooper JM, Hardy J, Wood NW, et al. Glucocerebrosidase deficiency in substantia nigra of parkinson disease brains. Ann Neurol 2012; 72: 455–63.

Grey M, Dunning CJ, Gaspar R, Grey C, Brundin P, Sparr E, et al. Acceleration of alpha-synuclein aggregation by exosomes. J Biol Chem 2015; 290: 2969–82.

Gündner AL, Duran-Pacheco G, Zimmermann S, Ruf I, Moors T, Baumann K, et al. Path mediation analysis reveals GBA impacts Lewy body disease status by increasing α-synuclein levels. Neurobiol Dis 2019; 121: 205–13.

Hannun YA, Obeid LM. Sphingolipids and their metabolism in physiology and disease. Nat Rev Mol Cell Biol 2018; 19: 175–91.

Hellstrand E, Nowacka A, Topgaard D, Linse S, Sparr E. Membrane lipid co-aggregation with alpha-synuclein fibrils. PLoS One 2013; 8: e77235.

Herzog R, Schuhmann K, Schwudke D, Sampaio JL, Bornstein SR, Schroeder M, et al. LipidXplorer: a software for consensual cross-platform lipidomics. PLoS ONE 2012; 7: e29851.

Herzog R, Schwudke D, Schuhmann K, Sampaio JL, Bornstein SR, Schroeder M, et al. A novel informatics concept for high-throughput shotgun lipidomics based on the molecular fragmentation query language. Genome Biol 2011; 12: R8.

Huebecker M, Moloney EB, van der Spoel AC, Priestman DA, Isacson O, Hallett PJ, et al. Reduced sphingolipid hydrolase activities, substrate accumulation and ganglioside decline in Parkinson’s disease. Mol Neurodegener 2019; 14: 40.

Imgrund S, Hartmann D, Farwanah H, Eckhardt M, Sandhoff R, Degen J, et al. Adult ceramide synthase 2 (CERS2)-deficient mice exhibit myelin sheath defects, cerebellar degeneration, and hepatocarcinomas. J Biol Chem 2009; 284: 33549–60.

Jones EE, Zhang W, Zhao X, Quiason C, Dale S, Shahidi-Latham S, et al. Tissue Localization of Glycosphingolipid Accumulation in a Gaucher Disease Mouse Brain by LC-ESI-MS/MS and High-Resolution MALDI Imaging Mass Spectrometry. SLAS Discov 2017; 22: 1218–28.

Koivusalo M, Jansen M, Somerharju P, Ikonen E. Endocytic trafficking of sphingomyelin depends on its acyl chain length. Mol Biol Cell 2007; 18: 5113–23.

Kroesen B-J, Jacobs S, Pettus BJ, Sietsma H, Kok JW, Hannun YA, et al. BcR-induced apoptosis involves differential regulation of C16 and C24-ceramide formation and sphingolipid-dependent activation of the proteasome. J Biol Chem 2003; 278: 14723–31.

Liebisch G, Binder M, Schifferer R, Langmann T, Schulz B, Schmitz G. High throughput quantification of cholesterol and cholesteryl ester by electrospray ionization tandem mass spectrometry (ESI-MS/MS). Biochim Biophys Acta 2006; 1761: 121–8.

Lingwood D, Simons K. Lipid rafts as a membrane-organizing principle. Science 2010; 327: 46–50.

Lippincott-Schwartz J, Phair RD. Lipids and cholesterol as regulators of traffic in the endomembrane system. Annu Rev Biophys 2010; 39: 559–78.

Martinez Z, Zhu M, Han S, Fink AL. GM1 specifically interacts with alpha-synuclein and inhibits fibrillation. Biochemistry 2007; 46: 1868–77.

Mazzulli JR, Xu YH, Sun Y, Knight AL, McLean PJ, Caldwell GA, et al. Gaucher disease glucocerebrosidase and alpha-synuclein form a bidirectional pathogenic loop in synucleinopathies. Cell 2011; 146: 37–52.

McNeill A, Magalhaes J, Shen C, Chau K-Y, Hughes D, Mehta A, et al. Ambroxol improves lysosomal biochemistry in glucocerebrosidase mutation-linked Parkinson disease cells. Brain 2014; 137: 1481–95.

Mencarelli C, Martinez-Martinez P. Ceramide function in the brain: when a slight tilt is enough. Cell Mol Life Sci 2013; 70: 181–203.

Middleton ER, Rhoades E. Effects of curvature and composition on α-synuclein binding to lipid vesicles. Biophys J 2010; 99: 2279–88.

Migdalska-Richards A, Wegrzynowicz M, Rusconi R, Deangeli G, Di Monte DA, Spillantini MG, et al. The L444P GBA mutation enhances alpha-synuclein induced loss of nigral dopaminergic neurons in mice. Brain 2017; 140: 2706–21.

Moors TE, Paciotti S, Ingrassia A, Quadri M, Breedveld G, Tasegian A, et al. Characterization of Brain Lysosomal Activities in GBA-Related and Sporadic Parkinson’s Disease and Dementia with Lewy Bodies. Mol Neurobiol 2019; 56: 1344–55.

Mosbech M-B, Olsen ASB, Neess D, Ben-David O, Klitten LL, Larsen J, et al. Reduced ceramide synthase 2 activity causes progressive myoclonic epilepsy. Ann Clin Transl Neurol 2014; 1: 88–98.

Mullin S, Smith L, Lee K, D’Souza G, Woodgate P, Elflein J, et al. Ambroxol for the Treatment of Patients With Parkinson Disease With and Without Glucocerebrosidase Gene Mutations: A Nonrandomized, Noncontrolled Trial. JAMA Neurol 2020; 77: 427–34.

Murphy KE, Halliday GM. Glucocerebrosidase deficits in sporadic Parkinson disease. Autophagy 2014; 10: 1350–1.

Neumann J, Bras J, Deas E, O’Sullivan SS, Parkkinen L, Lachmann RH, et al. Glucocerebrosidase mutations in clinical and pathologically proven Parkinson’s disease. Brain 2009; 132: 1783–94.

Niemelä PS, Hyvönen MT, Vattulainen I. Influence of chain length and unsaturation on sphingomyelin bilayers. Biophys J 2006; 90: 851–63.

Nussbaum RL. Genetics of Synucleinopathies. Cold Spring Harb Perspect Med 2018; 8: a024109.

O’Brien JS, Sampson EL. Lipid composition of the normal human brain: gray matter, white matter, and myelin. J Lipid Res 1965; 6: 537–44.

Olsen ASB, Færgeman NJ. Sphingolipids: membrane microdomains in brain development, function and neurological diseases. Open Biol 2017; 7

Polymeropoulos MH, Lavedan C, Leroy E, Ide SE, Dehejia A, Dutra A, et al. Mutation in the α-Synuclein Gene Identified in Families with Parkinson’s Disease. Science 1997; 276: 2045–7.

Rocha EM, Smith GA, Park E, Cao H, Brown E, Hallett P, et al. Progressive decline of glucocerebrosidase in aging and Parkinson’s disease. Ann Clin Transl Neurol 2015; 2: 433–8.

Ryan E, Seehra G, Sharma P, Sidransky E. GBA-associated parkinsonism: new insights and therapeutic opportunities. Current Opinion in Neurology 2019; 32: 589–596.

Sampaio JL, Gerl MJ, Klose C, Ejsing CS, Beug H, Simons K, et al. Membrane lipidome of an epithelial cell line. Proc Natl Acad Sci USA 2011; 108: 1903–7.

Sanchez-Martinez A, Beavan M, Gegg ME, Chau K-Y, Whitworth AJ, Schapira AHV. Parkinson disease-linked GBA mutation effects reversed by molecular chaperones in human cell and fly models. Scientific Reports 2016; 6: 31380.

Sardi SP, Clarke J, Kinnecom C, Tamsett TJ, Li L, Stanek LM, et al. CNS expression of glucocerebrosidase corrects alpha-synuclein pathology and memory in a mouse model of Gaucher-related synucleinopathy. Proc Natl Acad Sci USA 2011; 108: 12101–6.

Sassa T, Suto S, Okayasu Y, Kihara A. A shift in sphingolipid composition from C24 to C16 increases susceptibility to apoptosis in HeLa cells. Biochim Biophys Acta 2012; 1821: 1031–7.

Schapira AHV. Glucocerebrosidase and Parkinson disease: Recent advances. Molecular and Cellular Neuroscience 2015; 66: 37–42.

Schapira AHV, Olanow CW, Greenamyre JT, Bezard E. Slowing of neurodegeneration in Parkinson’s disease and Huntington’s disease: future therapeutic perspectives. Lancet 2014; 384: 545–55.

Schöndorf DC, Aureli M, McAllister FE, Hindley CJ, Mayer F, Schmid B, et al. iPSC-derived neurons from GBA-associated Parkinson’s disease patients show autophagic defects and impaired calcium homeostasis. Nat Commun 2014; 5: 4028.

Sidransky E, Nalls MA, Aasly JO, Aharon-Peretz J, Annesi G, Barbosa ER, et al. Multicenter Analysis of Glucocerebrosidase Mutations in Parkinson’s Disease. The New England Journal of Medicine 2009: 11.

Sillence DJ, Puri V, Marks DL, Butters TD, Dwek RA, Pagano RE, et al. Glucosylceramide modulates membrane traffic along the endocytic pathway. J Lipid Res 2002; 43: 1837–45.

van Smeden J, Janssens M, Boiten WA, van Drongelen V, Furio L, Vreeken RJ, et al. Intercellular skin barrier lipid composition and organization in Netherton syndrome patients. J Invest Dermatol 2014; 134: 1238–45.

Spillantini MG, Schmidt ML, Lee VM, Trojanowski JQ, Jakes R, Goedert M. Alpha-synuclein in Lewy bodies. Nature 1997; 388: 839–40.

Sun Y, Quinn B, Witte DP, Grabowski GA. Gaucher disease mouse models: point mutations at the acid beta-glucosidase locus combined with low-level prosaposin expression lead to disease variants. J Lipid Res 2005; 46: 2102–13.

Surma MA, Herzog R, Vasilj A, Klose C, Christinat N, Morin-Rivron D, et al. An automated shotgun lipidomics platform for high throughput, comprehensive, and quantitative analysis of blood plasma intact lipids. Eur J Lipid Sci Technol 2015; 117: 1540–9.

Winder-Rhodes SE, Evans JR, Ban M, Mason SL, Williams-Gray CH, Foltynie T, et al. Glucocerebrosidase mutations influence the natural history of Parkinson’s disease in a community-based incident cohort. Brain 2013; 136: 392–9.

Xu YH, Sun Y, Ran H, Quinn B, Witte D, Grabowski GA. Accumulation and distribution of α-synuclein and ubiquitin in the CNS of Gaucher disease mouse models. Mol Genet Metab 2011; 102: 436–47.

Yang S, Gegg M, Chau D, Schapira A. Glucocerebrosidase activity, cathepsin D and monomeric α-synuclein interactions in a stem cell derived neuronal model of a PD associated GBA mutation. Neurobiology of Disease 2020; 134: 104620.

Yang SY, Beavan M, Chau KY, Taanman JW, Schapira AH. A Human Neural Crest Stem Cell-Derived Dopaminergic Neuronal Model Recapitulates Biochemical Abnormalities in GBA Mutation Carriers. Stem Cell Reports 2017; 8: 728–42.

Yap TL, Velayati A, Sidransky E, Lee JC. Membrane-bound α-synuclein interacts with glucocerebrosidase and inhibits enzyme activity. Mol Genet Metab 2013; 108: 56–64.

Zhang Y, Shu L, Sun Q, Zhou X, Pan H, Guo J, et al. Integrated Genetic Analysis of Racial Differences of Common GBA Variants in Parkinson’s Disease: A Meta-Analysis. Front Mol Neurosci 2018; 11:43.

Zhu M, Fink AL. Lipid binding inhibits alpha-synuclein fibril formation. J Biol Chem 2003; 278: 16873–7.

