## Supplementary Information for "Sphingolipid changes in Parkinson L444P GBA mutation fibroblasts promote α-synuclein aggregation"

Running title: GBA Parkinson fibroblast lipid profile

Céline Galvagnion[1,2*], Silvia Cerri[3], Anthony H.V. Schapira[4], Fabio Blandini[3,5], Donato A. Di Monte[1*]

[1] German Center for Neurodegenerative Diseases (DZNE), Venusberg-Campus 1, Building 99, 53127 Bonn, Germany

[2] Current address: Department of Drug Design and Pharmacology, Faculty of Health and Medical Sciences, University of Copenhagen, Universitetsparken 2, 2100, Copenhagen Ø, Denmark

[3] Laboratory of Cellular and Molecular Neurobiology, IRCCS Mondino Foundation, Pavia, Italy

[4] Department of Clinical and Movement Neurosciences, UCL Queen Square Institute of Neurology, London, UK

[5] Department of Brain and Behavioral Sciences, University of Pavia, Pavia, Italy

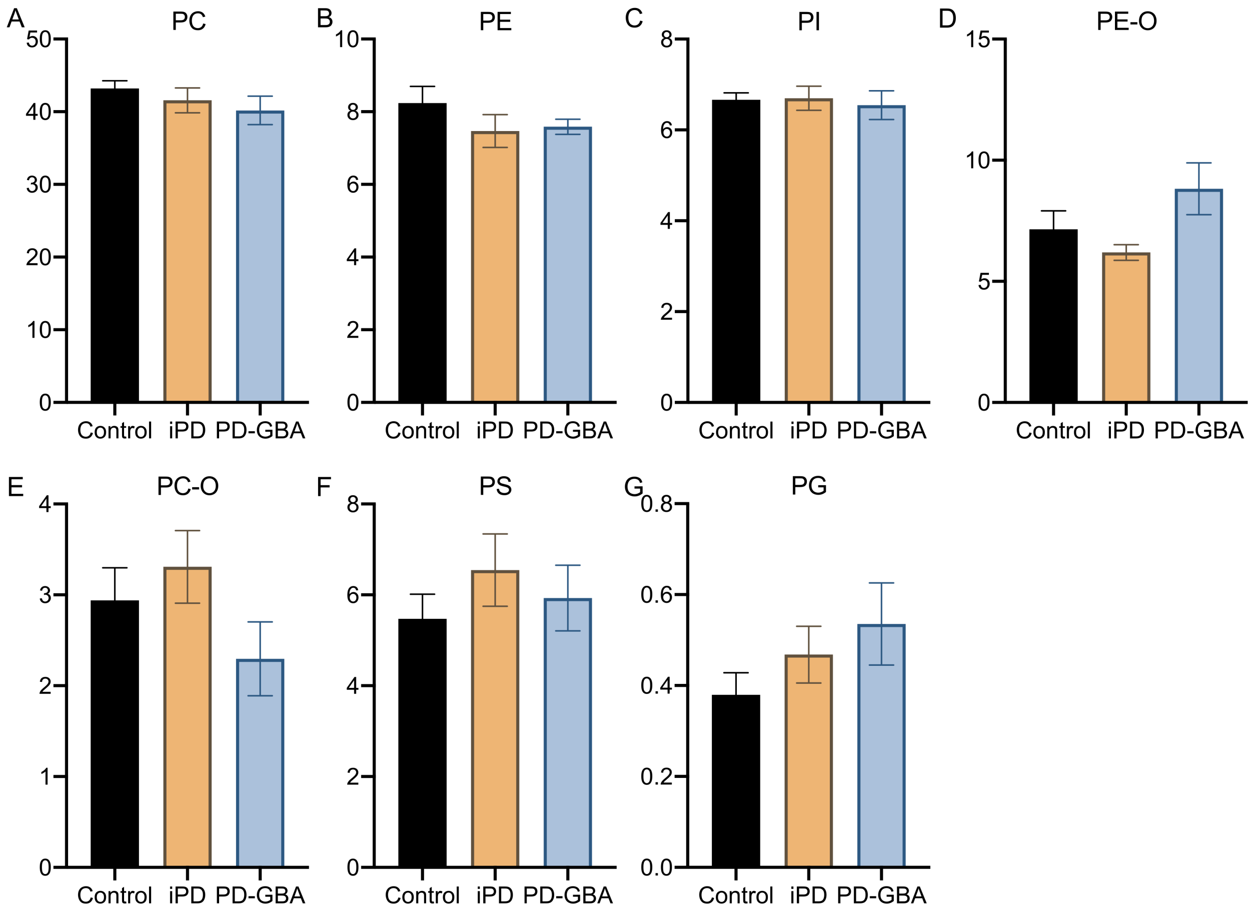


**Figure S1 Main species of phospholipids in control, iPD and PD-GBA fibroblasts.** Measurements of phosphatidylcholine (PC), phosphatidylethanolamine (PE), phosphatidylinositol (PI), PE ether (PE-O), PC ether (PC-O), phosphatidylserine (PS) and phosphatidylglycerol (PG) were made in fibroblast lipid extracts from control subjects ($n$ = 4) and iPD ($n$ = 4) and PD-GBA ($n$ = 5) patients. Data are expressed as percentage of the total lipid content. Bars show mean values, and error bars are ± SEM.

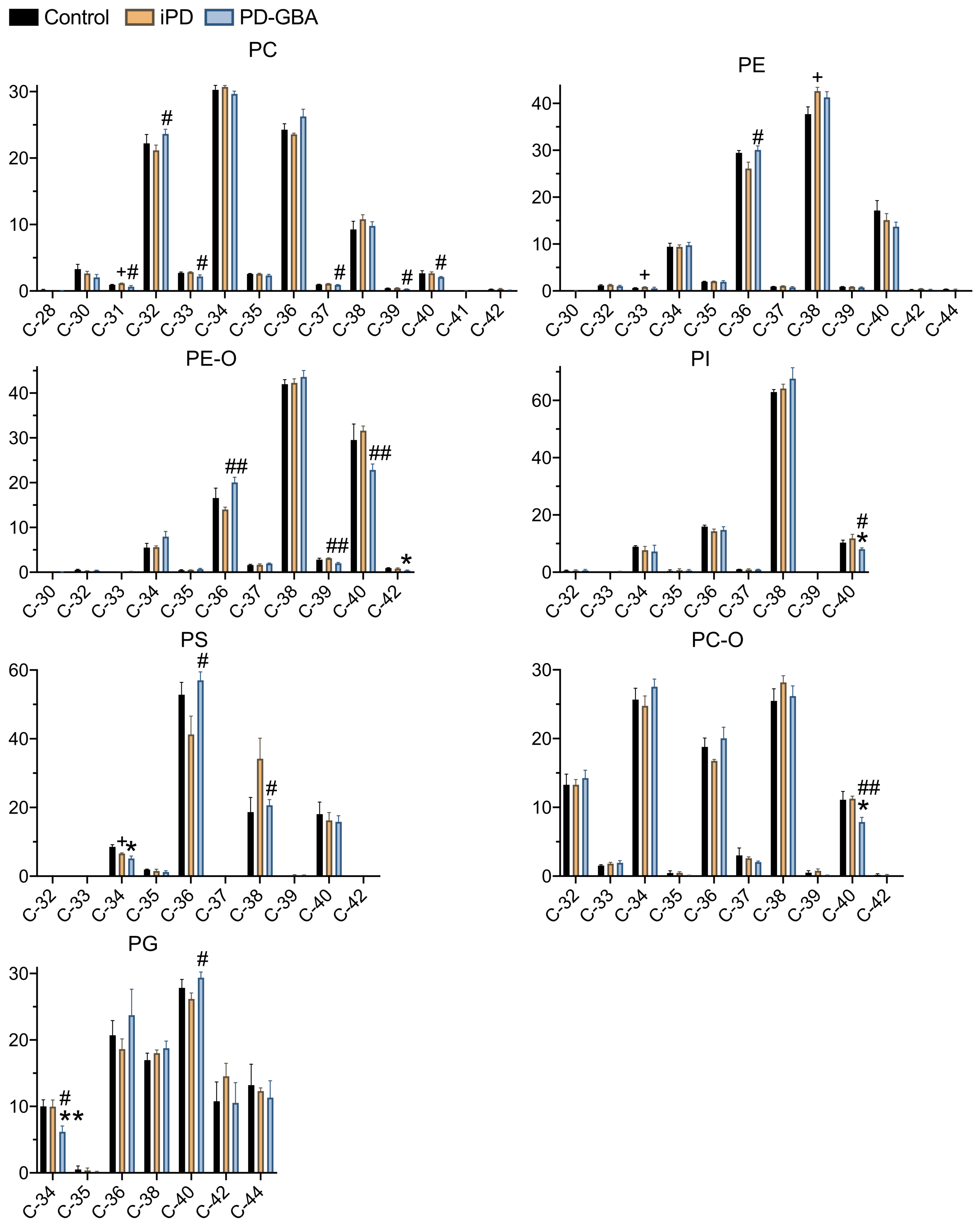


**Figure S2 Levels of phospholipid molecules with different acyl chain length.** Measurements of phosphatidylcholine (PC), phosphatidylethanolamine (PE), PE ether (PE-O), phosphatidylinositol

(PI), phosphatidylserine (PS), PC ether (PC-O) and phosphatidylglycerol (PG) with different hydrocarbon chain lengths (C-28 to C-44) were made in fibroblast lipid extracts from control subjects ($n$ = 4) and iPD ($n$ = 4) and PD-GBA ($n$ = 5) patients. Data are shown as percent of the respective total content. Bars show mean values, and error bars are ± SEM. Multiple $t$ test was performed to compare means between two groups, control *vs.* iPD ($^{+}$), control *vs.* PD-GBA (*) and iPD *vs.* PD-GBA ($^{\#}$). $^{+,*,\#}P < 0.05$; $^{**,\#\#}P < 0.005$. Control *vs.* PD-GBA (*,**): PE-O C-42 ($P$ = 0.0154), PI C-40 ($P$ = 0.0463), PS C-34 ($P$ = 0.0111), PC-O C-40 ($P$ = 0.0419), PG C-34 ($P$ = 0.0023). iPD *vs.* PD-GBA ($^{\#,\#\#}$): PC C-31 ($P$ = 0.0421), PC C-32 ($P$ = 0.0498), PC C-33 ($P$ = 0.0478), PC C-37 ($P$ = 0.0481), PC C-39 ($P$ = 0.0146), PC C-40 ($P$ = 0.0356), PE C-36 ($P$ = 0.0384), PE-O C-36 ($P$ = 0.0033), PE-O C-39 ($P$ = 0.0024), PE-O C-40 ($P$ = 0.0016), PI C-40 ($P$ = 0.0253), PS C-36 ($P$ = 0.0232), PS C-38 ($P$ = 0.0457), PC-O-C-40 ($P$ = 0.0046), PG C-34 ($P$ = 0.0257), PG C-40 ($P$ = 0.0365). Control *vs.* iPD ($^{+}$): PC C-31 ($P$ = 0.0438), PE C-33 ($P$ = 0.036), PE C-38 ($P$ = 0.0313), PS C-34 ($P$ = 0.0288).
